# Cellular adhesion is a controlling factor in neutrophil extracellular trap formation induced by antineutrophil cytoplasmic antibodies

**DOI:** 10.1101/2021.10.19.465054

**Authors:** Patrick M. Lelliott, Masayuki Nishide, Nicolas Pavillon, Yasutaka Okita, Takayuki Shibahara, Yumiko Mizuno, Hanako Yoshimura, Sho Obata, Atsushi Kumanogoh, Nicholas I. Smith

**Affiliations:** Laboratory of Biophotonics, Immunology Frontier Research Center, Osaka University, Osaka, Japan; Department of Respiratory Medicine and Clinical Immunology, Osaka University Graduate School of Medicine, Osaka, Japan; Department of Otorhinolaryngology-Head and Neck Surgery, Osaka University Graduate School of Medicine, Osaka, Japan; Laboratory of Immunopathology, Immunology Frontier Research Center, Osaka University, Osaka, Japan; Open and Transdisciplinary Research Institute (OTRI), Osaka University, Osaka, Japan

## Abstract

Anti-neutrophil cytoplasmic antibody (ANCA) associated vasculitis (AAV) is a life-threatening condition characterized by improper activation of neutrophils and release of neutrophil extracellular traps (NETs) in small vessels. This study aimed to explain the role of NETs in AAV pathogenesis by investigating a link between neutrophil adhesion and NET release. We leveraged an imaging flow cytometry-based assay and three-dimensional culture to demonstrate that neutrophil adhesion is essential for ANCA induced NET formation. We confirmed this requirement for cell adhesion using standard microscopy on ultra-low attachment hydrogel surfaces and demonstrate that this depends on the focal adhesion kinase pathway as determined using inhibitors for multiple targets in this process. ANCA increased expression of β2 integrins on neutrophils, and we confirmed that these integrins were required for NET formation using blocking antibodies. Finally, inhibitors for oxidative burst prevented NET formation, and this oxidative burst was mediated by the focal adhesion pathway. Overall, our findings reveal a central role for neutrophil attachment in NET formation in response to ANCA, helping to explain the restricted localization pattern of vessel damage, and suggesting that targeting neutrophil adhesion factors may be beneficial in preventing pathological damage from NETs during AAV.

## Introduction

Anti-neutrophil cytoplasmic antibodies (ANCA) occur during several forms of vasculitis, leading to the term ANCA-associated vasculitis (AAV) used for this collection of diseases^1^. AAV is characterized by damage to tissues and small vessels, particularly in the respiratory tract and kidneys, and can be clinically classified as granulomatosis with polyangiitis (GPA), microscopic polyangiitis (MPA), or eosinophilic granulomatosis with polyangiitis (EGPA). Although the prevalence of AAV is low, it has a significant impact on quality of life and mortality^2,3^, and a substantial economic burden^4^. ANCA consist of anti-myeloperoxidase (MPO) or anti-proteinase 3 (PR3) antibodies, these activate TNF-α Primed neutrophils, allowing them to adhere to endothelial cells^5^, degranulate^6,7^, and damage the endothelium^8^, with a range of pathological consequences in different tissues. More recently, the ability of neutrophils to form neutrophil extracellular traps (NETs) has been implicated in AAV pathogenesis^9^. This unique cellular response results in the unravelling of nuclear DNA, mixing of nuclear and cytoplasmic components, and finally release of large clouds of DNA peppered with various proteins such as histones and proteases^10^. While originally described as a potent weapon to immobilize and kill invading pathogens, NETs are now known to have a range of negative pathological effects, including damaging endothelial cells^11^. NETs may contribute to AAV directly through vascular damage, or, because ANCA antigens are abundant on NETs, may lead to antigen presentation and enhanced production of ANCA, creating a positive feedback loop^12^.

Although neutrophil attachment to the endothelium is known to play a critical part in AAV, the role of adhesion in ANCA induced NET formation has not been addressed. Indeed, studies investigating NETs almost exclusively involve the initial attachment of neutrophils to a surface to facilitate later analysis of NETs, without consideration of how attachment influences results. Exceptions to this include two recent studies. The first demonstrated that LPS induced NET formation requires substrate attachment, while PMA induced NETs do not^13^. The second study showed that substrate stiffness and coating dictate the ability of mouse-derived neutrophils to spontaneously form NETs^14^. These findings suggest that the control of neutrophil adhesion may offer a unique approach to mitigate pathological NET formation in diseases such as AAV. The therapeutic utility of targeting cell adhesion is increasingly being recognized, and adhesive molecules such as integrins and focal adhesion kinase (FAK) are being investigated as targets in a range of diseases and conditions such as cancer, inflammatory bowel disease, and multiple sclerosis, with some compounds already on the market and many others undergoing clinical trials^15–17^.

In this study, we investigated the influence of adhesion on the ability of neutrophils to form NETs in response to ANCA. We found that NET formation in response to ANCA, either commercially produced anti-MPO antibody or as total IgG isolated directly from AAV patient serum, was highly reliant on attachment, which could be blocked using antibodies for β2 integrins, or inhibitors of the focal adhesion pathway. We demonstrate the critical role of adhesion molecules in NET formation during AAV and suggest this can be exploited as a therapeutic pathway.

## Results

### NET formation in response to anti-MPO antibody stimulation requires substrate attachment

To investigate the role of adhesion in NET formation in ANCA vasculitis we stimulated neutrophils using anti-MPO antibodies. Peripheral blood neutrophils from healthy donors were stimulated with TNF-α and allowed to settle on either fibronectin-coated or commercially manufactured ultra-low attachment surface. The latter surface consists of a covalently bonded hydrogel proven to prevent neutrophil attachment, and we confirmed reduced neutrophil spreading on this surface with brightfield imaging (Figure 1A). Anti-MPO antibodies readily induced NETs on the fibronectin surface, while almost no NET formation occurred for neutrophils on the low attachment surface (Figure 1B). This was not the case for PMA stimulation, which induced NETs in both conditions. We tested collagen-coated, ICAM-1 coated, and uncoated tissue culture plastic, which resulted in similar levels of NET formation, indicating this effect is not limited to fibronectin (Supp Fig 1).

**Figure 1.**
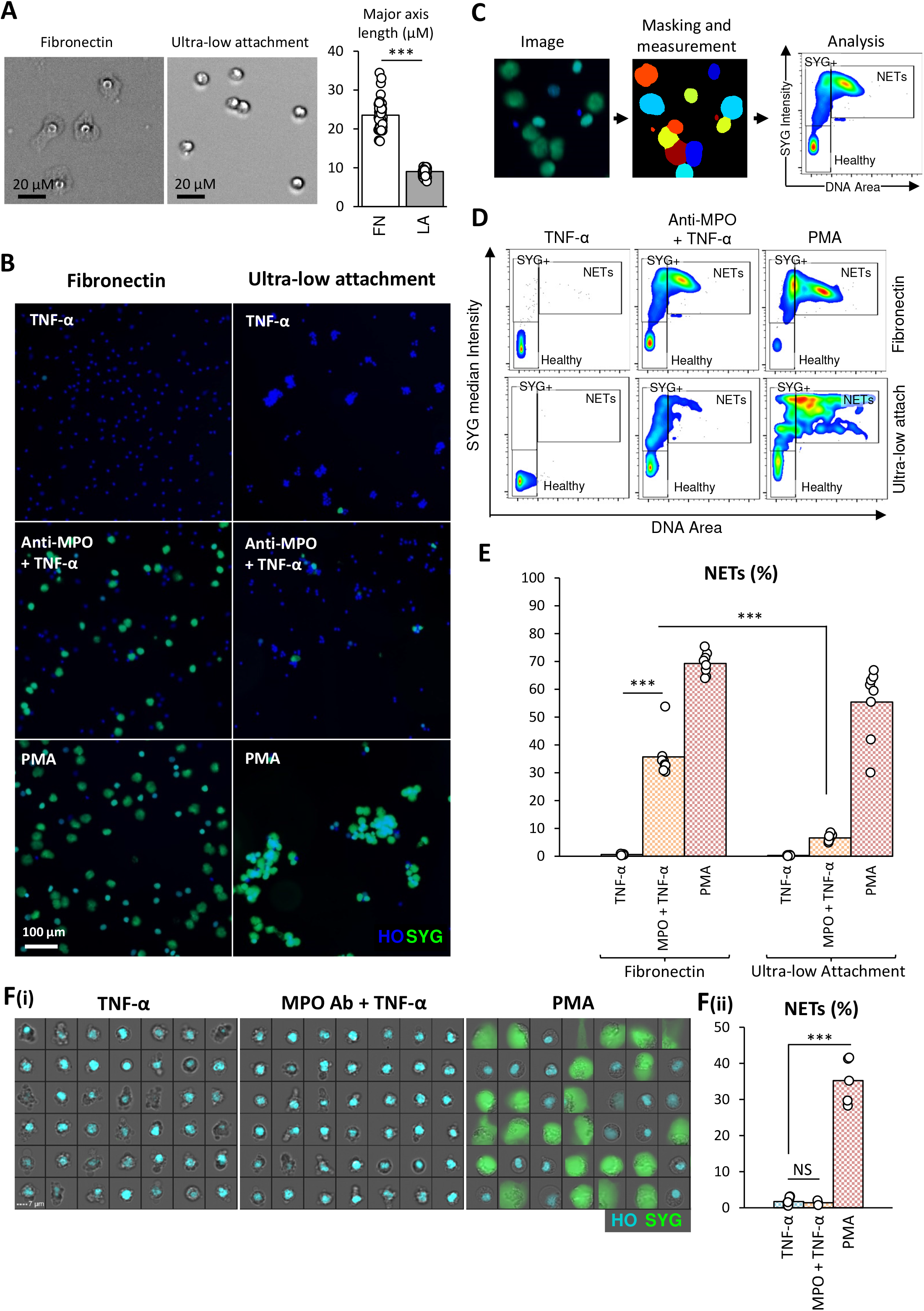
Substrate attachment is required for NET formation in response to anti-MPO antibody. Example images and major axis length of neutrophils on a fibronectin-coated (FN) versus an ultra-low attachment surface (LA) (A). Merged images of cells stained with Hoechst (blue) and Sytox Green (green) after treatment with TNF-α (5 ng/ml), and then left unstimulated, or stimulated with anti-MPO antibody (5 μg/mL) or PMA (100nM), and incubated for 4hrs in 96-well plates with the indicated surface (B). Masking of well images and measurement of area/shape and intensity using a CellProfiler pipeline, followed by analysis using FlowJo (C). NETs were quantified based on masked object area (DNA area) and median intensity of Sytox Green staining for each object. Representative plots are shown from one replicate for each condition (D). Percentage NETs for each condition with a minimum of 1000 cells analyzed for each well (E). Random example images and quantification of NETs measured by imaging flow cytometry using a previously published method^18^ (F). Data points represent technical replicates from one of three independent experiments with different donors. MPO – anti-MPO antibody, SYG – Sytox Green, HO – Hoechst 33342. NS = not significant, *** *p* < 0.001.

To quantify NET formation, we measured two parameters, DNA area and intensity of Sytox Green staining (SYG intensity) (Figure 1C). We plotted these two parameters and defined NETs as the DNA area (high) and SYG intensity (high) population. The SYG intensity (high) and DNA area (low) population were excluded from NET quantification as these cells more closely resemble necrotic cells without the release of DNA beyond the cell membrane. Approximately 36% of anti-MPO stimulated neutrophils formed NETs when incubated on a fibronectin-coated surface, while only 7% formed NETs on the low attachment surface (Figure 1D-E). For comparison, we found that neutrophils formed 69% and 55% NETs on these surfaces when stimulated with PMA. To further investigate the role of adhesion, we stimulated neutrophils in solution and analysed them by imaging flow cytometry using a method we have described previously^18^. Under these conditions, no NET formation could be detected with anti-MPO stimulation, while PMA stimulation-induced abundant NETs as expected (Figure 1F).

### Inhibition of the focal adhesion pathway prevents anti-MPO induced NET formation

Given the apparent importance of cell attachment in anti-MPO induced NET formation, we next investigated the focal adhesion signalling pathway. In neutrophils pre-incubated with FAK inhibitor-1, NET formation reverted to baseline levels (Figure 2A-C). FAK activation is facilitated by phosphorylation by Src kinases. As expected, inhibiting these kinases with Src inhibitor 1 and PP1 also reduced NET formation significantly, although PP1 inhibition was more effective. Actin polymerization is initiated by focal adhesions and allows for cell spreading and migration, as well as signalling at the focal adhesion junction through mechanical forces on FAK and Talin. Activation of these enzymes has widespread effects on cell processes, with a major target being PI3K. Inhibition of actin polymerization by cytochalasin B, and inhibiting PI3K both completely prevented NET formation in response to anti-MPO. Overall, these results indicate a critical role for the focal adhesion pathway in MPO induced NET formation.

**Figure 2.**
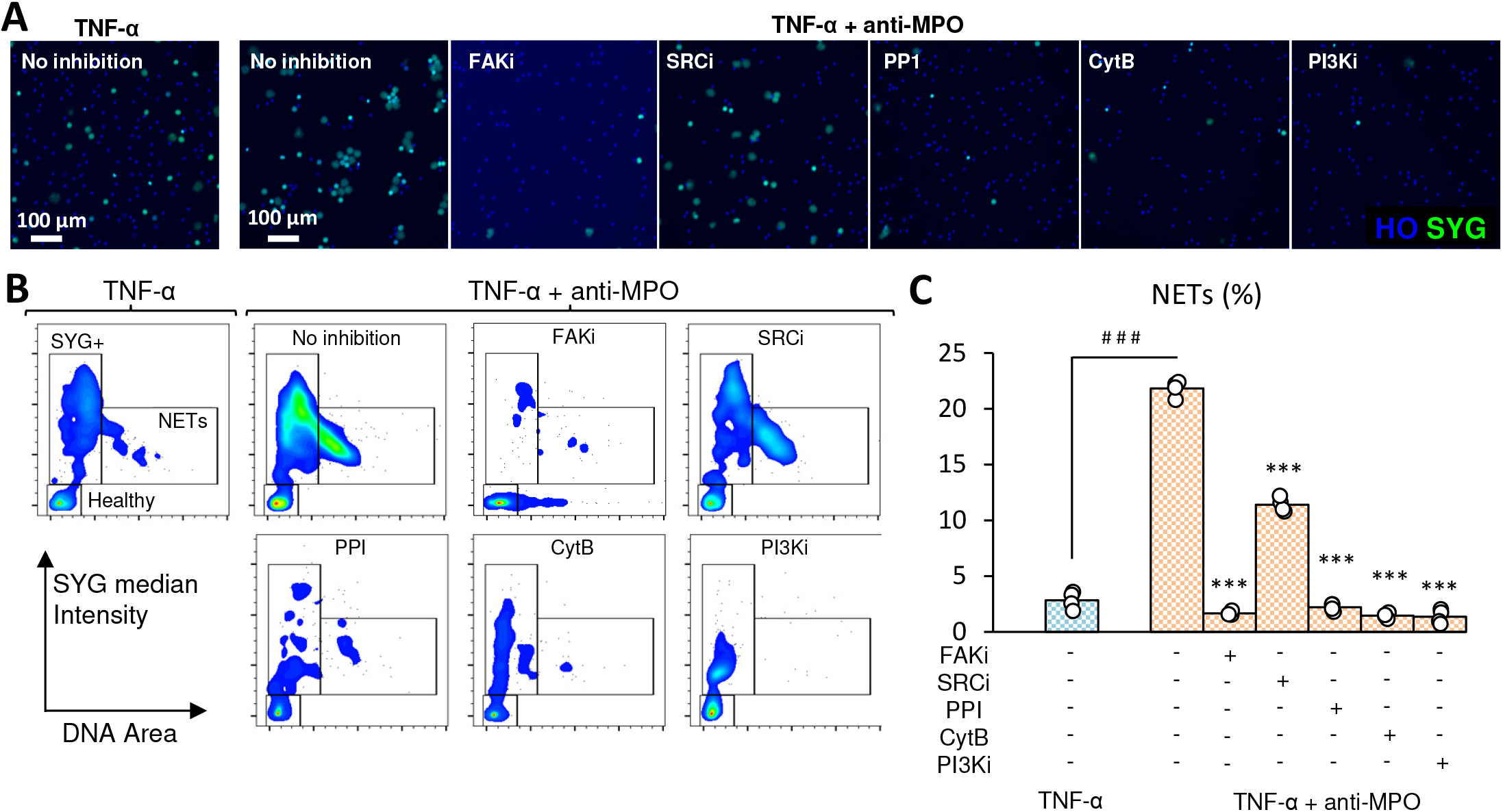
Inhibition of the focal adhesion pathway prevents anti-MPO induced NET formation. Merged images of peripheral blood human neutrophils stained with Hoechst (blue) and Sytox Green (green) after stimulation for 4hrs with TNF-α (5 ng/ml) and anti-MPO antibody (5 μg/mL) on fibronectin-coated plates with or without 30 minutes pre-incubation with inhibitors (A). Representative FlowJo plots of CellProfiler cell measurements (B), and percentage NETs (C) for each condition. Data points represent technical replicates from one of three independent experiments with different donors. MPO – anti-MPO antibody, SYG – Sytox Green, HO – Hoechst 33342, FAKi – Focal adhesion kinase inhibitor (20 μM), SRCi – Src kinase inhibitor-1 (1 μM), PP1 – Src kinase inhibitor (2.5 μM), CytB – actin filament inhibitor cytochalasin B (10 μM), PI3Ki – PI3K inhibitor copanlisib (200 nM). # # # *p* < 0.001 between mock and anti-MPO stimulated, *** *p* < 0.001 between no inhibition and with respective inhibitor.

### Involvement of ROS in NET formation

To provide a link between the focal adhesion pathway and NET formation, we considered the role of NADPH oxidase and ROS. Neutrophils incubated with the ROS scavenger pyrocatechol, and the NADPH oxidase inhibitor DPI, failed to produce NETs in response to anti-MPO (Figure 3A-B), indicating a ROS-dependant pathway of NET formation. The MEK-ERK pathway can be activated through PI3K and the focal adhesion pathway to phosphorylate and activate NADPH oxidase^19^. Consistent with this, the MEK inhibitor U0126 blocked NET formation. Downstream of ROS, NE is a critical enzyme involved in NET formation, and inhibiting this enzyme also blocked NET formation. We next measured ROS release in the context of anti-MPO stimulation with or without these inhibitors. ROS increased significantly with TNF-α and anti-MPO stimulation but remained at close to baseline levels with TNF-α alone (Figure 3C). Importantly, inhibition of FAK almost completely prevented ROS release, even under anti-MPO stimulation, indicating a combination of anti-MPO stimulation and activation of the focal adhesion pathway and is required before ROS production occurs. Downstream of FAK, Src kinase and MEK inhibition partially reduced ROS, while PI3K inhibition completely blocked ROS release. Inhibition of NADPH oxidase also prevented ROS release as expected. NE is activation is thought to occur downstream of ROS production. Consistent with this, NE inhibition had little effect on ROS release. Unexpectedly, inhibition of actin polymerisation did not reduce but rather increased ROS production. Together these results indicate a pathway in which attachment primes cells for ROS production leading to activation of NE and eventually NET release.

**Figure 3.**
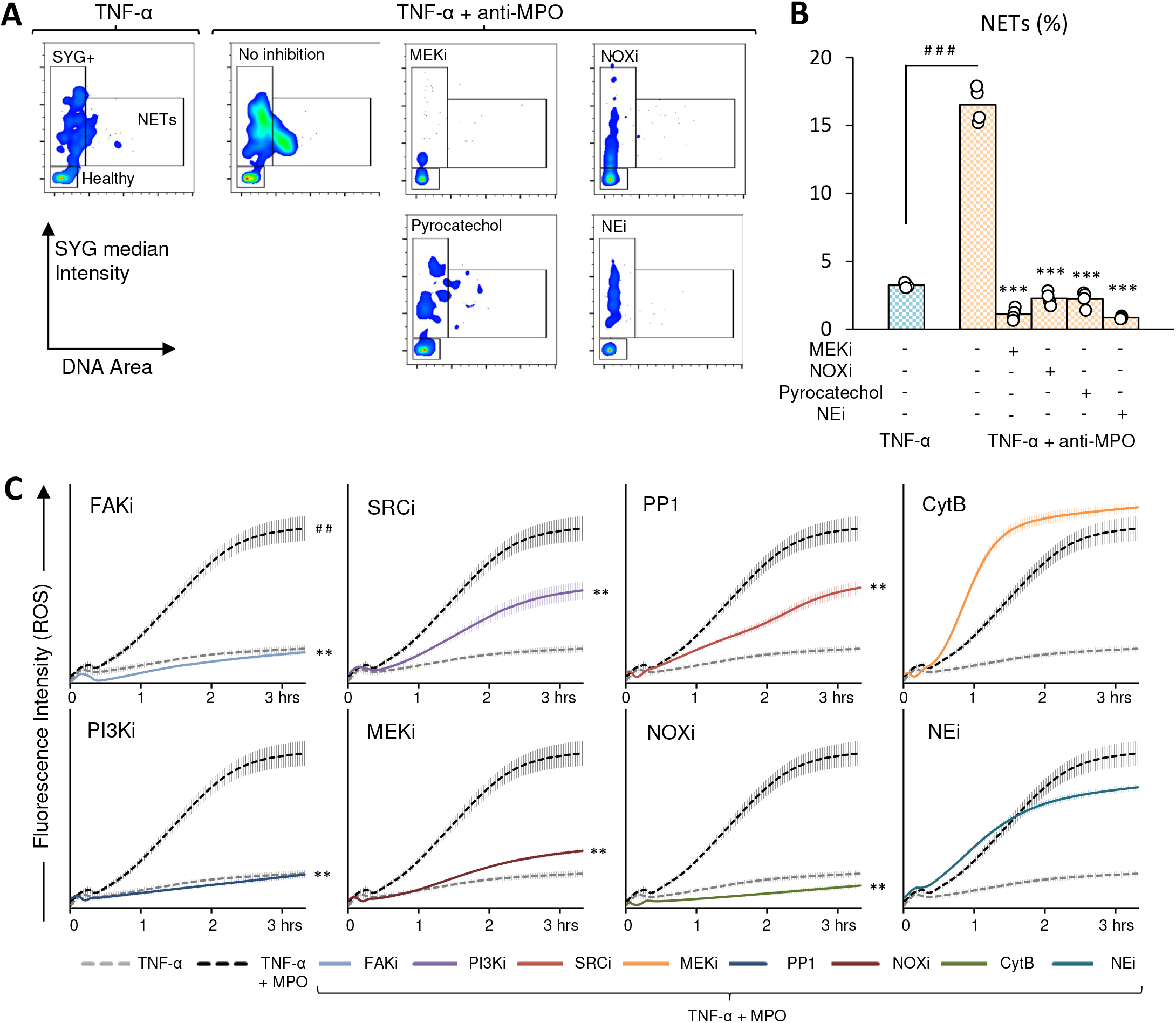
Involvement of ROS in anti-MPO induced NETs. Representative FlowJo plots of CellProfiler cell measurements (A), and percentage NETs (B) of neutrophils stimulated for 4hrs with TNF-α (5 ng/ml) and anti-MPO antibody (5 μg/mL) on fibronectin-coated plates with or without 30 minutes pre-incubation with inhibitors. Data points represent technical replicates from one of three independent experiments with different donors. Measurement of ROS release using a fluorometric kit monitored every two minutes over three hours. Mean and SEM is shown for four technical replicates. Data for each inhibitor is shown on a separate plot for clarity, with the TNF-α and anti-MPO stimulated results replicated from the same data. Significance is indicated for the final time point. MPO – anti-MPO antibody, FAKi – Focal adhesion kinase inhibitor (20 μM), SRCi – Src kinase inhibitor-1 (1 μM), PP1 – Src kinase inhibitor (2.5 μM), CytB – actin filament inhibitor cytochalasin B (10 μM), PI3Ki – PI3K inhibitor copanlisib (200 nM), MEKi – MEK inhibitor U0126 (25 μM), NOXi – NADPH oxidase inhibitor DPI (10 μM), Pyrocatechol – ROS scavenger (100 μM), NEi – neutrophil elastase inhibitor GW 311616A (20 μM). # # # *p* < 0.001, # # *p* < 0.01 between TNF-α and TNF-α + anti-MPO stimulated, *** *p* < 0.001, ** *p* < 0.01 between no inhibition and with respective inhibitor.

### Blocking of integrin β2 but not other major neutrophil integrins prevents anti-MPO NET formation

We next investigated which neutrophil adhesion molecules could be responsible for attachment and activation of the focal adhesion pathway using blocking antibodies. We focussed on integrin molecules known to be expressed on human neutrophils, integrin β1, β2, β3, α5, α9, αL, αM, and αX. Of all the integrins we tested, only blocking integrin β2 reduced NET formation significantly (Figure 4A-B), indicating the engagement of this integrin is critical for NET formation in response to anti-MPO antibodies. Furthermore, the expression of integrin β2 was significantly increased with TNF-α priming and anti-MPO stimulation, and synergistically in combination, as indicated by mean fluorescence intensity of neutrophils stained with anti-integrin β2 measured by flow cytometry (Figure 4Ci-ii). We next tested if β2 receptor ligation and crosslinking could prime neutrophils for NET formation, negating the need for cell attachment. However, neutrophils incubated on low attachment plates with crosslinking of β1, β2, β3, α5, α9, αL, αM, αX, or a combination of all, remained unresponsive to anti-MPO in terms of NET formation, indicating additional factors such as engagement of other receptors, cell spreading, or mechanical force signalling are required (Supp Fig 2).

**Figure 4.**
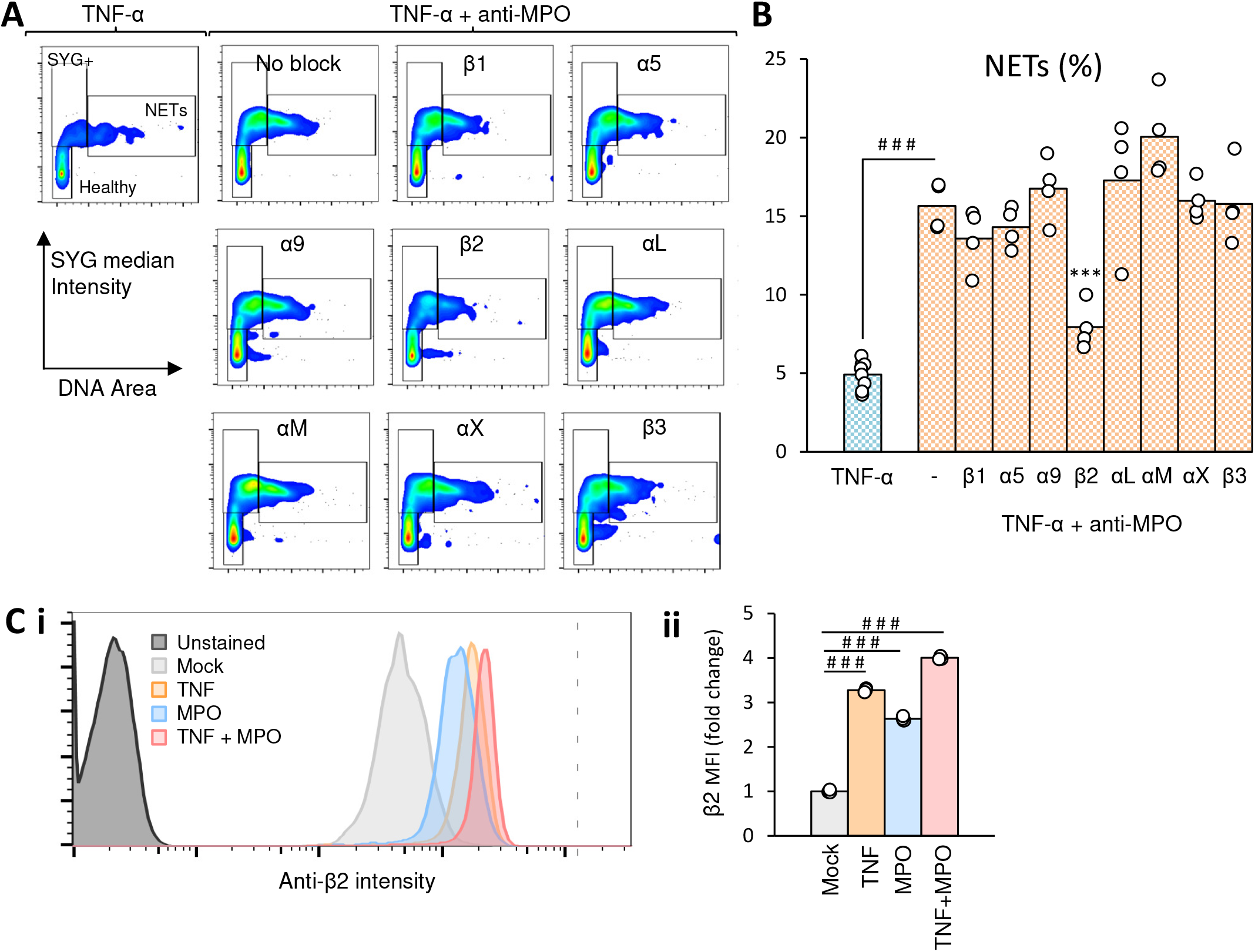
Formation of NETs in response to anti-MPO stimulation requires engagement of integrin β2. Peripheral blood human neutrophils were pre-incubated with TNF-α (5 ng/ml) and blocking antibodies for β1, β2, β3, α5, α9, αL, αM, or αX integrin (all 10 μg/ml) for 30 minutes with regular mixing to keep cells in suspension before being placed onto fibronectin-coated plates and stimulated with anti-MPO antibodies (5 μg/mL). Representative FlowJo plots of CellProfiler cell measurements (A), and percentage NETs (B) for each condition. Anti-β2 intensity histogram (C i), and mean fluorescence intensity (MFI) (C ii), of unstimulated (Mock), or neutrophils stimulated with TNF-α alone (5 ng/ml), anti-MPO alone (5 μg/mL), or TNF-α plus anti-MPO antibodies. Data points represent technical replicates from one of three independent experiments with different donors. # # # *p* < 0.001 between mock and stimulated, *** *p* < 0.001 between no blocking and with respective blocking antibody.

### AAV patient serum IgG induces attachment and focal adhesion-dependent NET formation

To confirm that our results with anti-MPO antibody stimulation could be reproduced using more clinically relevant stimuli, we collected serum from patients with ANCA vasculitis, who had not yet received immunosuppressive treatments, and isolated total IgG antibodies. Neutrophils stimulated with patient IgG induced significant NET formation, while IgG isolated from healthy controls did not (Figure 5A-C). In line with our results using anti-MPO antibodies, NET formation was highly dependent on cell adhesion, with almost no NETs present on the low attachment surface. NET formation was not correlated with clinical parameters such as BVAS or serum levels of MPO-ANCA, but it was positively correlated with the serum levels of C-reactive protein (CRP), suggested that the capacity of inducing NETs is associated with the severity of systemic inflammation (Supp Fig 3). Consistent with results using commercial anti-MPO antibodies, ANCA vasculitis patient sourced IgG induced NETs were prevented using inhibitors for FAK, Src kinase, actin polymerisation, and PI3K (Figure 5A-C). Overall, our results highlight a critical role for neutrophil attachment mediated by β2 integrins, the focal adhesion pathway, and ROS in the formation of NETs in response to ANCA, summarized in Figure 6.

**Figure 5.**
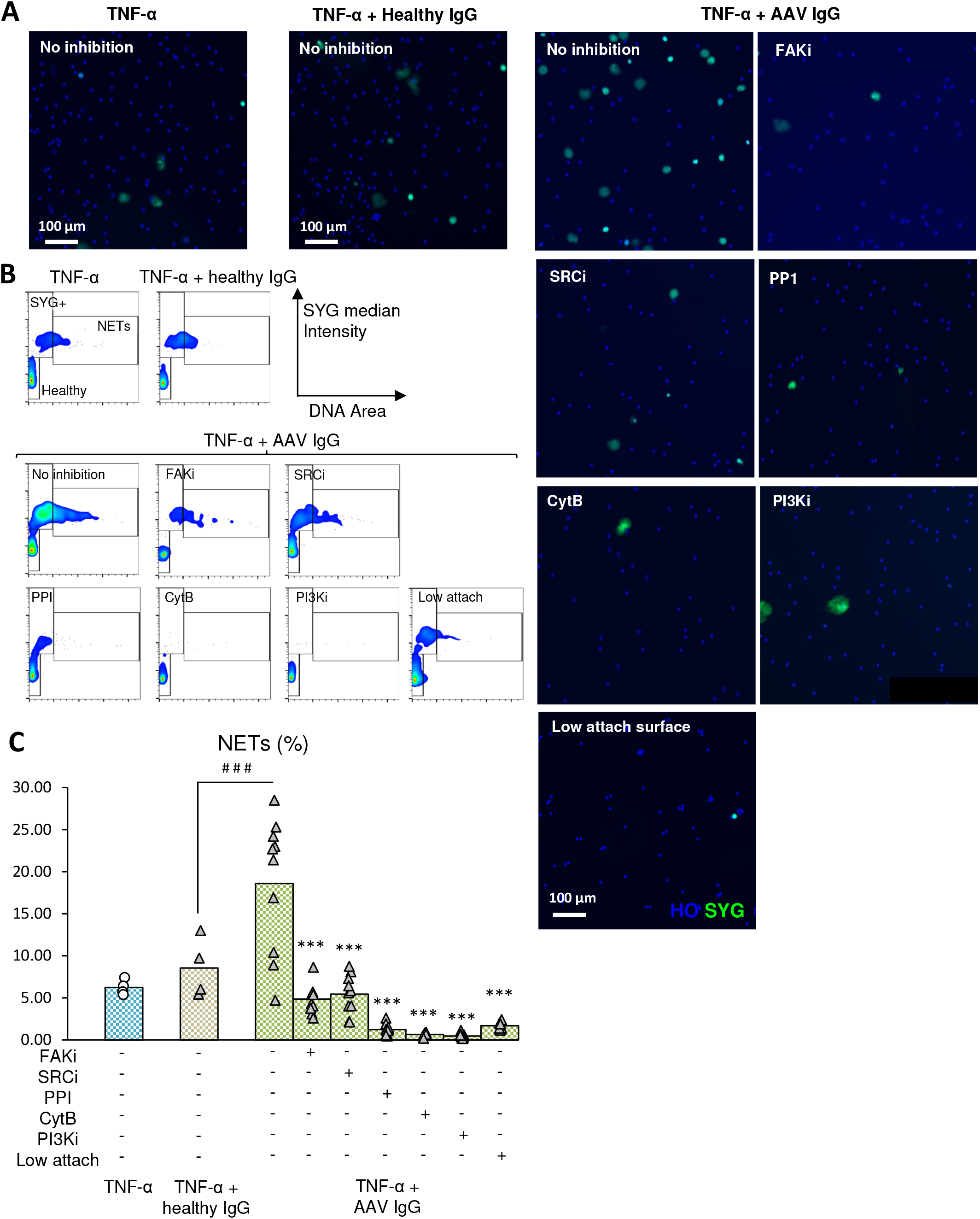
Immunoglobulins isolated from patients with ANCA vasculitis induce focal adhesion pathway dependent NET formation. Merged images of peripheral blood human neutrophils stained with Hoechst (blue) and Sytox Green (green) after stimulation for 4hrs with TNF-α (5 ng/ml) and isolated serum IgG (40 μg/mL) from ANCA vasculitis patients (N = 10) or healthy controls (N = 4), with or without 30 minutes pre-incubation with inhibitors (A). Representative plots of CellProfiler cell measurements (B), and percentage NETs (C) for each condition. For serum IgG, data points represent each patient or healthy control. For mock, data points represent technical replicates. Results are from one of three independent experiments with different healthy neutrophil donors. SYG – Sytox Green, HO – Hoechst 33342, FAKi – Focal adhesion kinase inhibitor (20 μM), SRCi – Src kinase inhibitor-1 (1 μM), PP1 – Src kinase inhibitor (2.5 μM), CytB – actin filament inhibitor cytochalasin B (10 μM), PI3Ki – PI3K inhibitor copanlisib (200 nM). # # *p* < 0.01 between healthy and ANCA serum IgG, *** *p* < 0.001 between no inhibition and with respective inhibitor.

**Figure 6.**
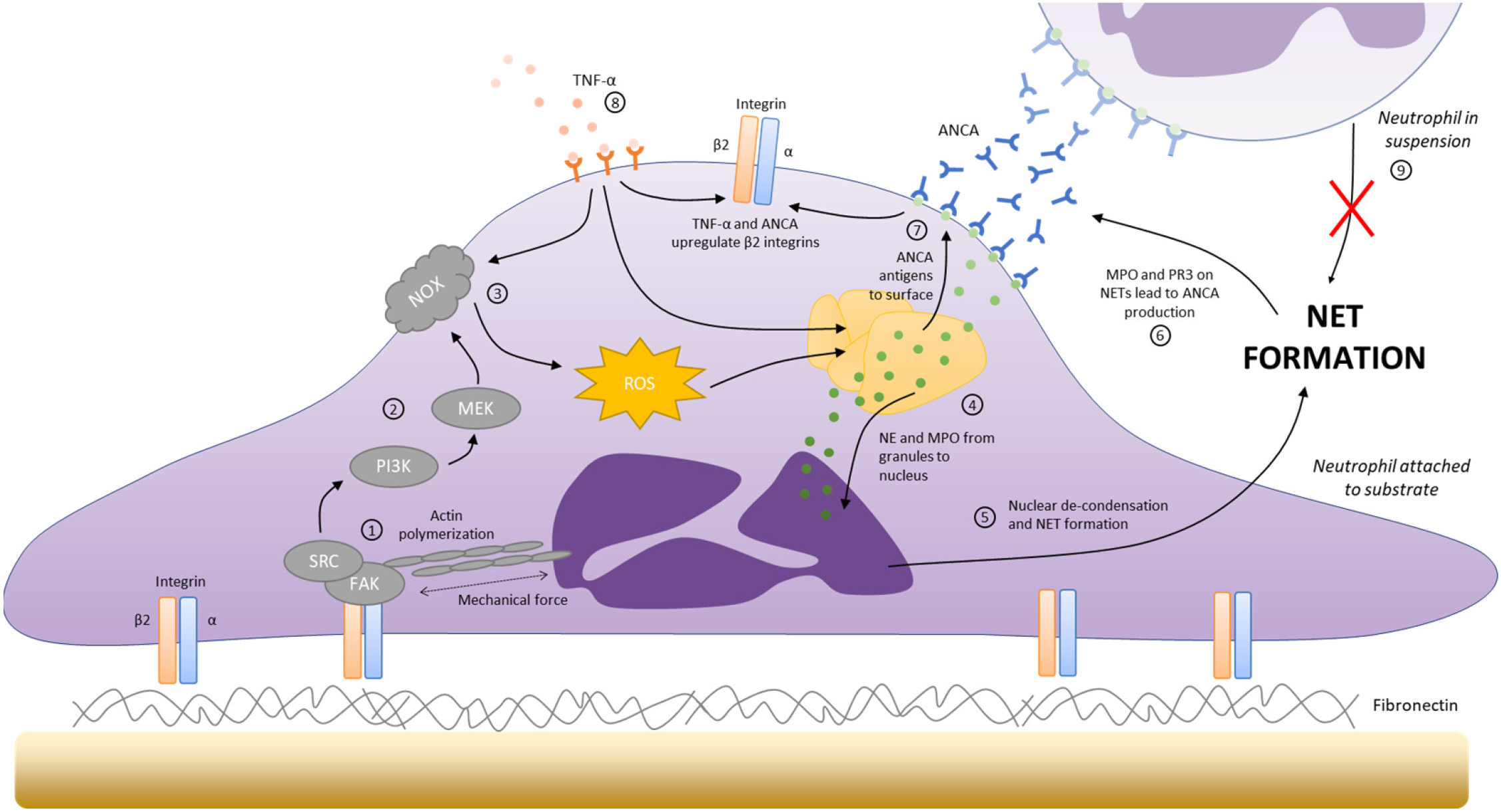
Proposed model for the involvement of adhesion in ANCA induced NET formation. Engagement of β2 integrins with fibronectin initiates formation of the focal adhesion complex involving Src, FAK, and actin polymerization (1)^36–38^ (this study). PI3K and MEK are activated downstream of focal adhesions (2)^42^, which leads to ROS production by NADPH oxidase (NOX) (3)^67^ (this study). ROS leads to the translocation of NE and MPO to the nucleus (4)^48^, which starts the process of DNA de-condensation, nuclear membrane rupture, mixing of cytoplasmic contents and DNA, and finally breakdown of the cell membrane and NET release (5)^48^. ANCA antigens present on NETs encourage the formation of ANCA and the breakdown of immune tolerance (6)^12^. ANCA interact with antigens on the neutrophil surface, leading to increased β2 integrins and ROS^57^ (this study), creating a positive feedback loop (7). TNF-α contributes to this process in several ways, encouraging the activation of NOX to produce ROS^68^, increasing the expression of β2 integrins^58^ (this study), and increasing membrane levels of ANCA antigens^7^ (8). Neutrophils in suspension do form NETs in response to ANCA (9) (this study).

## Discussion

This study reveals an underappreciated role for neutrophil attachment in dictating NET formation in response to ANCA both in the form of commercial anti-MPO antibodies and isolated IgG from AAV patients. NET formation was mediated by the focal adhesion pathway and ROS production, and required integrin β2, with other major neutrophil integrins being dispensable.

Neutrophil adhesion is known to play an important role in the pathology of AAV^20–22^. Inflammatory cytokines such as TNF-α are known pathological factors during AAV^23,24^, and along with complement C5a and ANCA, these reduce neutrophil deformability^25,26^ and encourage neutrophil aggregation^27^, thereby promoting interactions with the vessel walls, particularly in the small capillaries which are the main site of AAV associated pathology^28^. While neutrophil responses to adhesion such as ROS release and degranulation have been well-studied^29^, there are comparatively few studies that have addressed the role of adhesion in NET formation. One study found that adhesion was required for LPS induced NET formation but was dispensable when cells were stimulated with PMA^13^, while a second study found that substrate stiffness correlated with spontaneous NET formation in mice^14^. In the case of AAV, the role of adhesion in NET formation was previously unknown. Preventing neutrophil adhesion and measuring NET formation in suspension is extremely difficult due to the delicate nature of NETs. While several conventional flow cytometry approaches claim to identify NETs in suspension, these have not attempted to distinguish NETs from other types of cell death^30–32^, and their ability to measure NETs has been questioned^33^. More advanced imaging flow cytometry techniques have also claimed to identify NETs^34,35^, however, these failed to show the clouds or strings of DNA typically considered NETs. None of these studies addressed potential differences in NET formation when adhered versus in suspension. We recently developed a protocol which maintains NETs in suspension using density control, followed by analysis using imaging flow cytometry, and demonstrated the ability to image fully formed NETs identified as large clouds of DNA^18^. Here, we used this approach to demonstrate that unlike PMA induced NET formation, ANCA induced NET formation is completely blocked in the absence of adhesion. We further corroborated this using ultra-low attachment surfaces, on which neutrophils also failed to produce NETs in response to ANCA.

Engagement of integrins during neutrophil attachment initiates the focal adhesion pathway. Central to this pathway is FAK, whose activation is mediated through phosphorylation by Src kinase^36–38^. In our study, we demonstrate that both FAK and Src are required for ROS production and NET formation. In the case of Src, we used two different inhibitors, Src inhibitor 1 and PP1, providing 40% and 38% reduction in ROS, and 48% and 90% reduction in NETs respectively. PP1 is known to cause partial off-target inhibition of p38 MAPKs^39^, which are reportedly involved in NET formation in response to PMA^40^. This dual-target inhibition of PP1 may explain its increased effect on NET formation compared with the more specific Src inhibitor 1. PI3K activation occurs downstream of FAK^41,42^, and we show that inhibitors of PI3K also block both ROS and NET formation. Although PI3K is a central kinase involved in many cellular processes, this finding nevertheless adds weight to the importance of the FAK pathway in NET formation and may help explain previously reported results for the involvement of PI3K in NET formation, such as in response to parasites, microcrystals, and PMA^43,44^. An important consequence of integrin engagement and FAK activation is cell remodelling and actin polymerization. Actin provides a link to the focal adhesion site, allowing the cell to move and stretch, actions which can be detected through force-sensitive proteins at the focal adhesion junction such as Talin^45^. The lack of NET formation in neutrophils in which we crosslinked integrin receptors to mimic adhesion conditions provides evidence for the importance of mechanical signalling, but our finding that inhibition of actin polymerisation does not prevent ROS production suggests that this process is more important in later stages of NET formation, although this requires further study.

ROS production is a clear candidate linking cell attachment to NET formation. ROS has been implicated in many pathways of NET formation, most commonly through NADPH oxidase activity^46^. Indeed, this was the case in our study, with NETs completely dependent on both ROS and NADPH oxidase. There are a large variety of pathways that can activate NADPH oxidase, including many pathways initiated through focal adhesion, and it is well known that neutrophil ROS activity is strongly influenced by adhesion. In this study, we focused on MEK, an enzyme recently reported to be required for PMA induced NET formation^47^, as it is known to be activated downstream of PI3K. This proved to be the case for our study too, with ANCA induced ROS largely prevented, and NET formation completely blocked by MEK inhibition. Finally, NADPH oxidase ROS production leads to release and activation of granule proteins such as NE, which is well known to be a critical enzyme involved in NET formation^48,49^, and we found was also required in our study.

Inflammation or damage to the endothelium triggers expression of adhesion molecules such as ICAM-1 and deposition of plasma fibronectin^50–52^, which provide attachment sites for normally benign neutrophils^53,54^. Integrins, particularly β2 integrins such as MAC-1 (αMβ2) and LFA-1 (αLβ2), provide attachment points to these sites^55,56^. Both ANCA and TNF-α are known to increase expression of β2 integrins on the neutrophil surface^57,58^, and this was the case in our study. It is plausible that a combination of cytokine priming and neutrophil attachment is required to overcome inhibitory mechanisms and result in NETs in response to ANCA. In support of this, we found when we blocked β2 integrins, ANCA induced NET formation was reduced to nearly background levels. This is consistent with a previous study reporting that β2 integrin is required for neutrophil respiratory burst in response to ANCA^59^. In a clinical setting, patients with active AAV show increased expression of both β1 and β2 integrins^60^. Although β1 integrins have been reported to enhance neutrophil attachment to endothelial cells, specifically α5β1^61^ and α9β1^62^, in our case we found no effect of blocking these. Likewise, we found no evidence for the involvement of β3 integrins. It should be emphasized that our study was limited in examining only the interaction of β2 integrins with fibronectin. The situation *in vivo* is considerably more complex, with numerous integrin-ligand binding interactions occurring depending on the local microenvironment. Indeed, we found attachment to a wide range of surfaces was sufficient to allow NET formation in response to ANCA.

Overall, our results help to clarify the complex feedback cycle of neutrophil activation in AAV. Low-level neutrophil activation induced by TNF-α and ANCA results in increased surface β2 integrins which promote neutrophil aggregation^57,58 63^, encouraging physical trapping of aggregates in capillaries. Neutrophil arrest in small capillaries is enhanced by adhesion mediated by endothelial receptor expression, and deposition of microparticles and plasma fibronectin^51,52,64^. Adhesion of neutrophils further promotes activation and endothelial inflammation through degranulation and oxidative burst^29^, and here we demonstrate it is also required for NET formation in the presence of ANCA. NET formation contributes to this feedback cycle in two ways. NETs contain ANCA antigens and have been shown to enhance their immunogenicity, resulting in increased ANCA production^12^. NETs have also been shown to damage endothelial cells^11^, which would enhance the deposition of plasma fibronectin and increase the expression of endothelial receptors. Thus, our study postulates a central role of neutrophil adhesion in ANCA pathogenesis and highlights the potential utility of therapies aimed at controlling or preventing pathological neutrophil adhesion.

## Methods

### Human subjects

Human samples were obtained after informed consent was provided in accordance with the Declaration of Helsinki and with approval from the ethical review board of the Graduate School of Medicine, Osaka University, Japan (no. T19204 and no. 11122-5). Patients with AAV were diagnosed as having microscopic polyangiitis (MPA) or granulomatosis with polyangiitis (GPA), according to the criteria of the Research Committees of the Japanese Ministry of Health, Labour and Welfare. GPA was also diagnosed according to the American College of Rheumatology classification criteria. Disease activity score of AAV was estimated by Birmingham Vasculitis Activity Score (BVAS ver3). IgG was isolated from healthy control or AAV patient serum using a Melon™ Gel IgG Spin Purification Kit (Thermo Fisher Scientific, Waltham, MA).

### Neutrophil isolation

To mitigate neutrophil adherence during preparation we performed isolation within 30 minutes of blood collection, used low-attachment labware, and kept isolated neutrophils for a maximum of 1 hour before use, maintaining them in suspension. Neutrophils were isolated from blood using an EasySep Direct Human Neutrophil Isolation Kit (Stemcell Technologies, Vancouver, Canada), with 0.05% human serum albumin (HSA, Sigma-Aldrich, St. Louis, MO) added to the isolation buffer. Isolated neutrophils were washed in Hanks Buffered Salt Solution (HBSS, Thermo Fisher Scientific) with 0.05% HSA.

### Measurement of neutrophil extracellular trap formation

For fluorescence microscopy, neutrophils in Dulbecco’s Modified Eagle Medium: Nutrient Mixture F-12 (DMEM/F12) with 15 mM HEPES and 0.05% HSA, without phenol red (Thermo Fisher Scientific) were added to either tissue culture-treated plates coated with 10 μg/ml human plasma fibronectin (Sigma-Aldrich) or Corning® Costar® Ultra-Low Attachment plates (Sigma-Aldrich), which are coated with a proprietary hydrogel designed to minimise cell attachment. Cells were stimulated with 5 ng/ml TNF-α (Peprotech, Rocky Hill, NJ) followed by 5 μg/ml anti-MPO antibodies (A0398 polyclonal, Agilent Technologies, Santa Clara, CA) or 40 μg/ml isolated serum IgG, and incubated for 4 hours at 37 °C with 5% CO_2_. In some cases, cells were stimulated with 100 nM phorbol 12-myristate 13-acetate (PMA) without TNF-α. Cells were stained with 4 μM Hoechst 33342 (Sigma-Aldrich) as a membrane-permeable DNA stain, and 500 nM Sytox Green (Thermo Fisher Scientific) as a membrane-impermeable DNA stain, and imaged without washing steps using either a Toxinsight (Thermo Fisher Scientific) or CQ1 (Yokogawa, Tokyo, Japan) fluorescence microscope with 10X objective. In some cases, neutrophils were pre-incubated with 20 μM focal adhesion kinase (FAK) inhibitor-1 (Merck, Darmstadt, Germany), 1 μM Src inhibitor-1 (Merck), 2.5 μM Src inhibitor PP1 (Cayman Chemical, Ann Arbor, MI), 10 μM actin inhibitor cytochalasin B from Helminthosporium dematioideum (Nacalai Tesque, Kyoto, Japan), 200nM phosphoinositide 3-kinase (PI3K) inhibitor copanlisib (Cayman Chemical), 25 μM of the MEK inhibitor U0126 (Nacalai Tesque), 100 μM of the ROS scavenger pyrocatechol (Sigma-Aldrich), 10 μM of the NADPH oxidase inhibitor DPI (Sigma-Aldrich), 20 μM of the NE inhibitor GW 311616A (Axon Medchem, Netherlands), or 10 μg/ml azide-free blocking antibodies for integrin β1 (CD29, clone: TS2/16), β2 (CD18, clone: TS1/18), β3 (CD61, clone: 23C6), α5 (CD49e, clone: NKI-SAM-1), αL (CD11a, clone: HI111), αM (CD11b, clone: M1/70), αX (CD11c, clone: 3.9) (all Biolegend, San Diego, CA), or α9 (CD49i, clone: Y9A2, Merck) for 30 minutes at 37 °C before stimulation, with gentle agitation every 10 minutes to prevent cell settling.

Imaging flow cytometry of NETs was performed as previously described^18^. Briefly, neutrophils in DMEM/F12 with 15 mM HEPES and 0.05% HSA, without phenol red, and with 1.6 μM Hoechst 33342 (Sigma-Aldrich), and 25 nM SYTOX Green (Thermo Fisher Scientific), were incubated for 4 hours at 37 °C with 5% CO_2_ with stimulants in low attachment 1.5 mL tubes (PROKEEP, Watson Biolab, Kobe, Japan) with caps open. 10 μL 90% v/v Percoll in DMEM/F12 was layered under cells during incubation to minimise cell aggregation and interactions with the tube walls and create a three-dimensional culture environment for the cells. After incubation cells were fixed in 1% paraformaldehyde (PFA) (Sigma-Aldrich) for 10 minutes at room temperature, then gently resuspended and analysed as-is on an ImageStream X Mark II instrument (Luminex, Austin, TX) using a 60X objective. Events with brightfield area or SYTOX green area greater than 20 μm^2^ were collected. Compensation was performed using single stained controls.

### Image analysis and neutrophil extracellular trap quantification

Fluorescent microscopy images were analysed using Cellprofiler V4.0.6^65^ using a custom pipeline. Hoechst stains all DNA, while Sytox Green will be excluded from healthy cells and will only stain DNA within dead cells, or DNA present in NETs. Sytox Green staining was brighter than Hoechst staining, which was especially important for defining diffuse NET structures. Therefore, these images were combined, using each pixel’s maximum value from the two channels, creating a greyscale image of merged fluorescence that represents the overall distribution of DNA. This image was used to identify objects using the Otsu thresholding approach. Sytox Green staining intensity and area have been previously used to classify healthy cells and NETs, and here we used a similar approach^18^. We measured two parameters for each object. The object area, defined as the number of pixels in the object, which we refer to as ‘DNA area’, and Sytox Green median intensity, which is obtained by overlaying the object masks onto the original Sytox Green image and measuring the median pixel intensity for each object. Object measurements were exported into one CSV file for each well of the experiment. CSV files were converted to FCS format, and Flowjo V10.7.1 was used for further analysis. DNA area was plotted against Sytox Green median intensity for each object, and three populations were defined, DNA Area (low) Sytox Green (low) - Healthy neutrophils, DNA Area (low) Sytox Green (high) – Sytox Green positive neutrophils (non-NETs), Area (high) Sytox Green (high) – NETs.

Imaging flow cytometry images were processed using IDEAS v6.2 (Luminex) and FlowJo as described previously^18^. Briefly, the ‘Object(M01,Ch01,Tight) (Brightfield)’ and ‘Morphology(M02,Ch02) (Sytox Green)’ masks were used to identify cell and DNA boundaries, and the feature ‘Intensity_Morphology(M02,Ch02) And Not Object(M01,Ch01,Tight)_Ch02’ was calculated and defined as ‘exDNA (intensity)’. NETs were classified as the ‘exDNA (intensity)’ (high), ‘Intensity_MC_Ch07 (HO (Intensity))’ (high), population.

### ROS measurement

Isolated neutrophils were treated and prepared as described for NET measurements. ROS release was monitored for three hours with measurements every two minutes using a fluorometric hydrogen peroxide assay kit (Sigma-Aldrich) and a Glomax microplate reader (Promega, Madison, WI).

### Measurement of integrin expression

Isolated neutrophils were stimulated with 5 ng/ml TNF-α, 5 μg/ml anti-MPO antibodies, a combination of these, or left untreated. After 30 minutes incubation at 37 °C with gentle agitation every 10 minutes, Fc receptors were blocked with Human TruStain FcX™ (Biolegend), and cells were stained with 1.5 μg/ml Brilliant Violet 605™ anti-CD16 (clone: 3G8) and 1 μg/ml APC anti-CD18 (clone: 1B4/CD18) (Biolegend). Cell data was collected using an Attune flow cytometer (Thermo Fisher Scientific) and analysed with Flowjo. Single neutrophils were gated based on FSC/SSC and CD16 positive staining. The mean fluorescence intensity of CD18 on neutrophils was then measured.

### Statistics

Data were analysed by two-way ANOVA with interactions and post hoc Tukey tests using R v3.6.1^66^. A p-value < 0.05 was considered significant. Data are presented as the mean with individual data points or standard error indicated.

## Supporting information

All Supplemental Figures

## Acknowledgements

This work was funded by the Japan Society for the Promotion of Science (JSPS) through the Funding Program for World-Leading Innovative R&D on Science and Technology (FIRST Program); the JSPS World Premier International Research Centre Initiative Funding Program; the Uehara Memorial Foundation; the Centre of Innovation program (COISTREAM) from the Ministry of Education, Culture, Sports, Science and Technology of Japan (MEXT) (to A.K.); JSPS KAKENHI (JP18H05282 to A.K., JP18K16146 to M.N., JP19K23865 to P.M.L.); the Japan Agency for Medical Research and Development (AMED)(20ek0109307h0003, J200705023, J200705710, J200705049, JP18cm016335 and JP18cm059042 to A.K.); the Kansai Economic Federation (KANKEIREN); the Mitsubish Zaidan1 (to A.K.).

## Author contributions

PML and MN conceived of and designed the study. PML carried out the experiments. PML, MN, NP, AK, and NIS analysed and interpreted the data. YO, TS, YM, HY, SO contributed to preparation of materials and recruited and clinically characterised patients. PML wrote manuscript. MN, NP, AK, and NIS reviewed and edited the manuscript. All authors approved the final version of the manuscript.

