## Supplementary material for "Cellular adhesion is a controlling factor in neutrophil extracellular trap formation induced by antineutrophil cytoplasmic antibodies": All Supplemental Figures

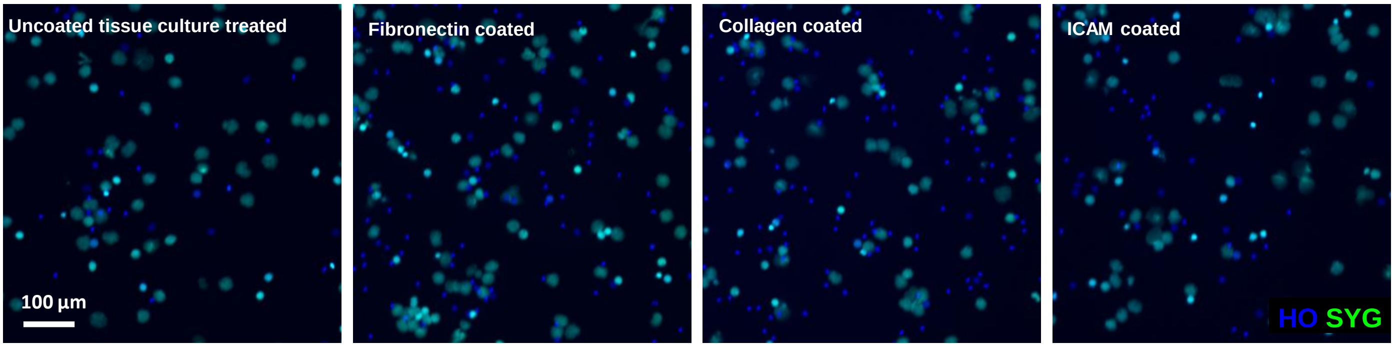


**Supplementary figure 1. Anti-MPO induced NETs on different substrates.** Merged images of peripheral blood human neutrophils stained with Hoechst (blue) and Sytox Green (green) after stimulation for 4hrs with TNF-α (5 ng/ml) and anti-MPO antibody (5 µg/mL) on plates with the indicated coatings. SYG – Sytox Green, HO – Hoechst 33342.

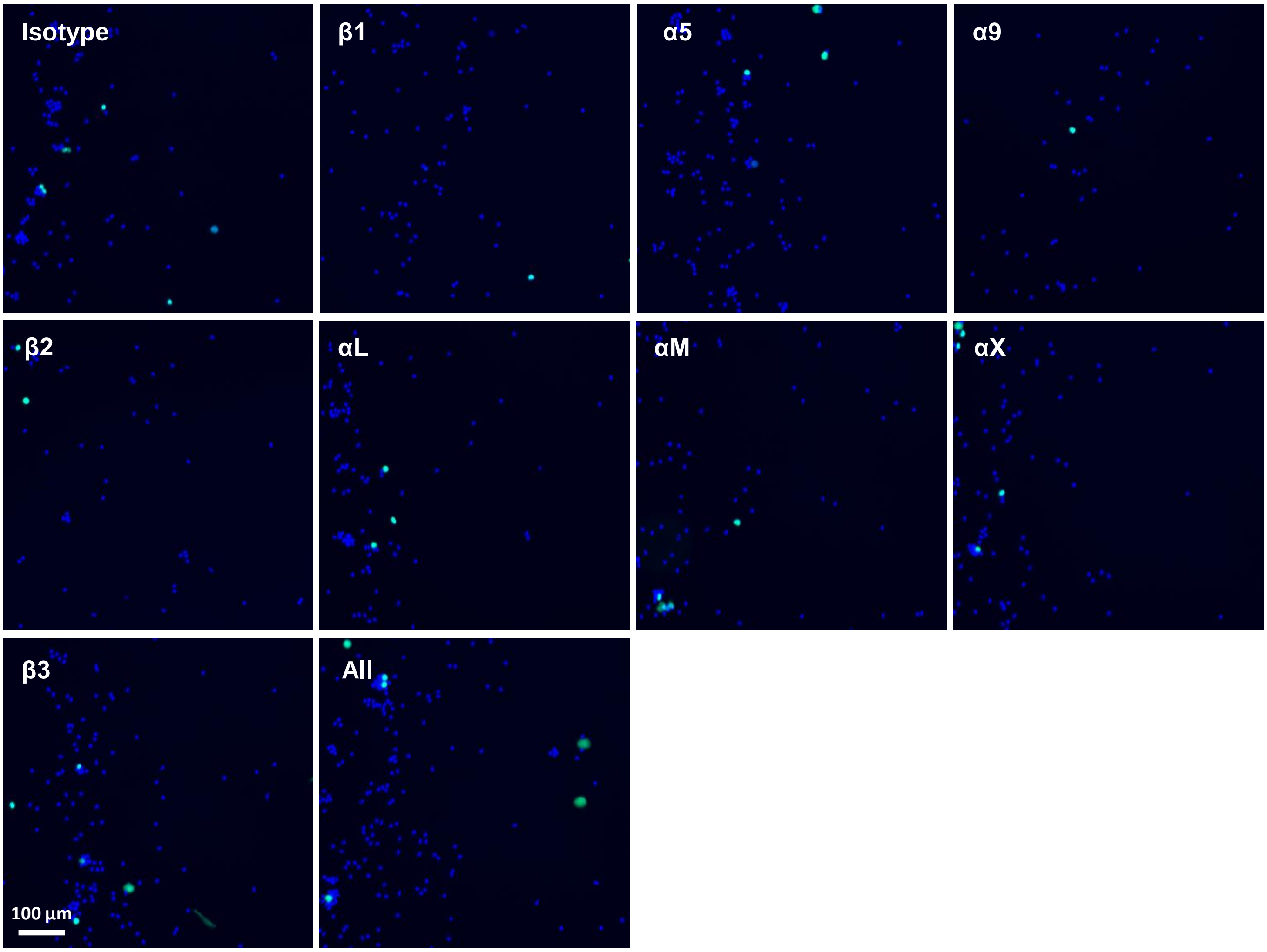


**Supplementary figure 2. Ligation and crosslinking of neutrophil integrins does not influence NET formation.** Merged images of peripheral blood human neutrophils stained with Hoechst (blue) and Sytox Green (green) after stimulation for 4hrs with TNF-α (5 ng/ml) and anti-MPO antibody (5 µg/mL) on ultra low attachment plates. Antibodies for the indicated integrin subunits were added and crosslinked using secondary antibodies to mimic integrin attachment during stimulation. All – combined crosslinking of all antibodies, SYG – Sytox Green, HO – Hoechst 33342.

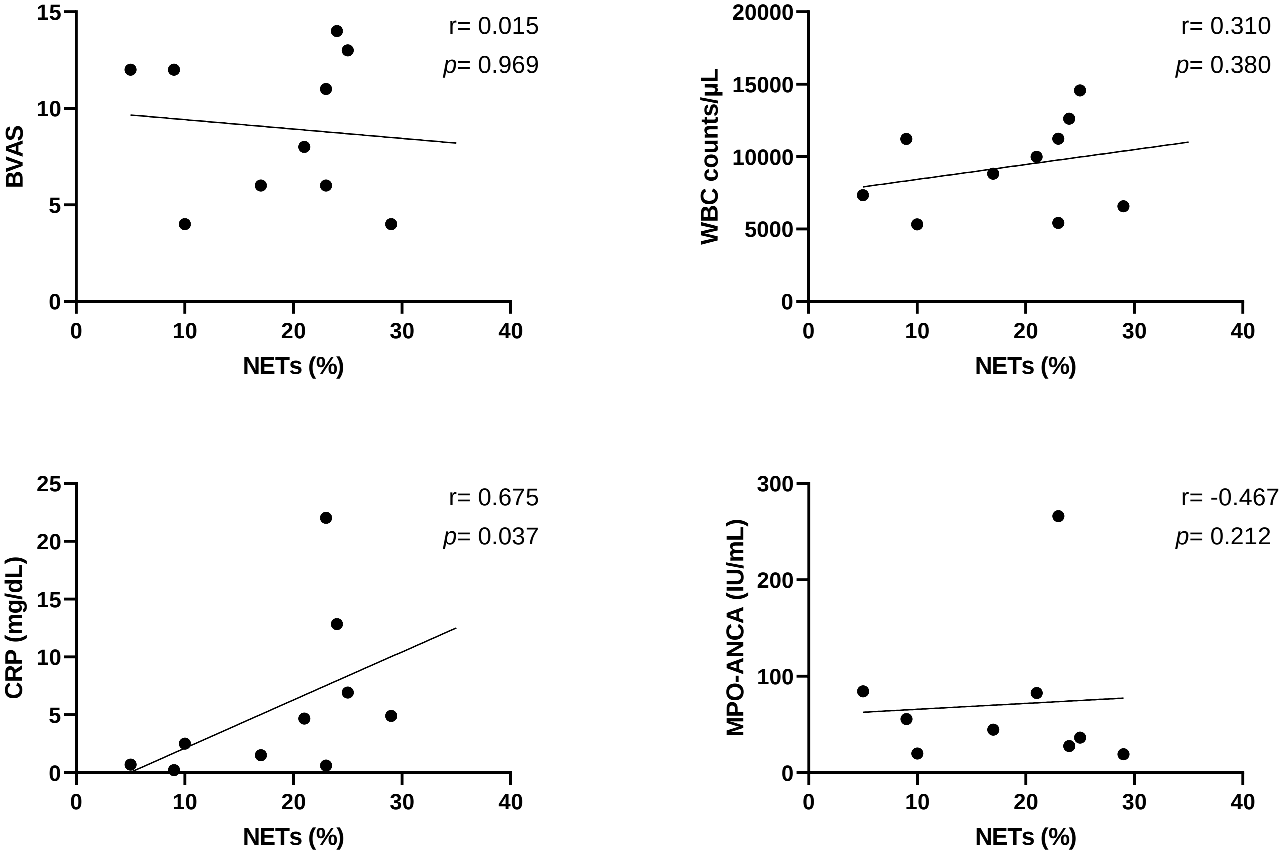


**Supplementary figure 3.**

Correlation of NETs formations by patients serum IgG with Birmingham Vasculitis Activity Score version 3 (BVAS), blood white blood cell counts, C-reactive protein (CRP) levels and MPO-ANCA titers. Correlations (r) and p values were quantified using Spearman’s rank correlation coefficient.
